# Single-cell RNA analysis reveals unexpected hemocyte plasticity and immune cell specialization in a *Drosophila* overgrowth model

**DOI:** 10.64898/2026.01.09.698745

**Authors:** Prathibha Yarikipati, Andreas Bergmann

## Abstract

Immune cells play essential roles in maintaining tissue homeostasis and responding to abnormal growth, but how innate immune cells adapt to chronic apoptotic signaling remains poorly understood. In *Drosophila melanogaster*, hemocytes, particularly plasmatocytes, are recruited to tumor-like overgrowths, yet their transcriptional diversity and lineage dynamics under these conditions remain undefined. Here, we apply single-cell RNA sequencing to nearly 50,000 circulating and sessile hemocytes from larvae bearing undead overgrown eye discs, a model of regenerative overgrowth driven by sustained caspase activity. We resolve 17 transcriptionally distinct hemocyte clusters, including known lineages and 13 previously unrecognized plasmatocyte subtypes. Interestingly, specific plasmatocyte populations are differentially expanded or depleted under overgrowth conditions. Notably, we identify a matrix-remodeling plasmatocyte population marked by high expression of Jonah-family serine proteases. Pseudotime analysis reveals unexpected plasmatocyte plasticity and two novel terminally differentiated effector states. These findings define the immune landscape of tumor-like overgrowth and establish *Drosophila* as a platform for dissecting innate immune responses to tissue stress and dysregulated growth *in vivo*.

## Introduction

The systemic immune response to tumorigenesis plays a fundamental role in determining disease outcome. In mammals, tumor-associated immune cells such as macrophages (TAMs) are deeply integrated into the tumor microenvironment (TME), where they can promote or suppress tumor growth through diverse functions, including immunosuppression, extracellular matrix remodeling, and angiogenesis (Kloosterman and Akkari, 2023; Ma et al., 2022; van Vlerken-Ysla et al., 2023). Dissecting the origins, transcriptional states, and functional plasticity of these immune cells has proven challenging due to the complexity of the mammalian immune system. As a genetically tractable model with a simplified innate immune system, *Drosophila melanogaster* provides a powerful platform for exploring conserved principles of immune-tumor interactions (Bilder et al., 2021).

In *Drosophila*, immune cells, collectively known as hemocytes, are generated via two hematopoietic waves. The first occurs in the embryo, giving rise to embryonic hemocytes that persist as circulating and sessile cells throughout larval life (Banerjee et al., 2019). The second wave takes place in the lymph gland, a larval hematopoietic organ that becomes active in third instar larvae and discharges hemocytes into circulation during pupation or immune challenge (Koranteng et al., 2022; Monticelli et al., 2024). Traditionally, hemocytes have been classified into three major lineages. Plasmatocytes constitute the majority (90-95%) of hemocytes and function as professional phagocytes. Crystal cells regulate melanization and wound healing through prophenoloxidase activity. Lamellocytes are large flattened cells that differentiate upon immune challenge and encapsulate pathogens such as parasitoid wasp eggs (Honti et al., 2014). More recently, thanocytes which are characterized by high expression of *Tep4* and *Ance*, and primocytes which express stem-like transcriptional regulators, have been identified through single-cell RNA sequencing (scRNA-seq) (Fu et al., 2020).

Recent scRNA-seq analyses have significantly advanced our understanding of *Drosophila* hematopoiesis and immune cell plasticity. Several seminal studies have mapped hemocyte transcriptional states under steady-state and immune-activated conditions. For example, Cattenoz et al. (2020) defined 14 distinct hemocyte clusters across developmental stages and in response to wasp infestation, identifying distinct subpopulations of plasmatocytes involved in phagocytosis, lipid metabolism, and stress response (Cattenoz and Giangrande, 2021; Cattenoz et al., 2021; Cattenoz et al., 2020). Tattikota et al. (2020) further resolved hemocyte heterogeneity into 17 clusters, and uncovered complex developmental trajectories and inter-hemocyte communication networks including an FGF signaling axis between crystal cells and lamellocytes (Tattikota et al., 2020). Fu et al. (2020) identified novel hemocyte types including thanocytes and primocytes (Fu et al., 2020). Additional studies further expanded on the hemocyte heterogeneity in *Drosophila* (Cho et al., 2020; Coates et al., 2021; Girard et al., 2021; Hultmark and Ando, 2022; Leitao et al., 2020; Mase et al., 2021; Shin et al., 2020) including pupal stages (Hirschhauser et al., 2023).

While these studies have already given a glimpse of the plasticity of *Drosophila* hemocytes, they have primarily focused on responses to infection or injury. In contrast, little is known about how hemocytes respond transcriptionally to chronic tissue stress, overgrowth and tumor-like conditions, where apoptotic and inflammatory signals persist. Several tumor models in *Drosophila* (*Ras^V12^scrib*, *Ras^V12^*, *vps25*) recruit hemocytes to overgrowing tumor tissues, with parallels to mammalian TAMs (Diwanji and Bergmann, 2020; Hirooka et al., 2025; Khalili et al., 2023; Pastor-Pareja et al., 2008; Perez et al., 2017; Voutyraki et al., 2023). However, the molecular identities, lineage trajectories, and functional states of hemocytes under these conditions remain poorly characterized.

A particularly relevant and genetically defined tumor-like context is apoptosis-induced proliferation (AiP) (Bergmann, 2025; Bergmann and Fan, 2025; Diwanji and Bergmann, 2019; Fogarty and Bergmann, 2017). In this process, cells that initiate apoptosis but are prevented from dying through expression of the effector caspase inhibitor *p35*, remain “undead” and paradoxically promote tissue overgrowth (Huh et al., 2004; Kondo et al., 2006; Perez-Garijo et al., 2004; Ryoo et al., 2004; Wells et al., 2006). These undead cells activate initiator caspases, generate extracellular reactive oxygen species (ROS), and release mitogens, leading to non-autonomous overproliferation of neighboring cells. When *hid* and *p35* are expressed in the eye imaginal disc (*ey>hid;p35*), the result is a robust, caspase-dependent overgrowth of the head tissue (Fan et al., 2014). Hemocytes are strongly recruited to these undead discs, attracted at least in part by ROS signaling (Fogarty et al., 2016). More recent work suggests that basement membrane (BM) damage in these tissues further contributes to hemocyte recruitment, resembling neoplastic models such as *Ras^V12^scrib* (Diwanji and Bergmann, 2020; Pastor-Pareja et al., 2008; Perez et al., 2017). However, hemocytes do not appear to directly cause BM damage, as there is no correlation between hemocyte density and BM degradation, suggesting instead that they are responding to localized tissue damage.

Despite these parallels to tumor biology, the diversity and transcriptional states of hemocytes in overgrowth models remain largely unexplored. Most notably, it is unclear whether these hemocytes adopt specialized effector programs, similar to TAMs in mammals, or represent generic inflammatory responses. A recent study analyzed hemocytes in a *Drosophila* tumor model (*Ras^V12^*) in salivary glands and identified tumor-associated transcriptional changes, including upregulation of stress and immune response genes (Khalili et al., 2023). However, the small cell numbers precluded the discovery of rare subtypes or intermediate cell states, and did not address lineage relationships among hemocytes.

To address these gaps, we performed high-throughput single-cell RNA sequencing of hemocytes from larvae expressing *ey>hid;p35* to investigate how caspase-induced tissue overgrowth shapes the systemic immune response. We profiled 49,039 hemocytes from undead and control conditions, including both circulating and sessile populations, and identified 17 transcriptionally distinct hemocyte clusters, including known types and a rich array of novel plasmatocyte subpopulations. Several of these clusters are selectively enriched in the AiP background, revealing context-dependent immune states. In particular, we identify a matrix-remodeling plasmatocyte cluster marked by secreted serine proteases of the Jonah family, which are likely involved in basement membrane repair. We also reconstructed hemocyte developmental trajectories using Monocle3, and identified two previously unrecognized terminal plasmatocyte states. Therefore, by characterizing immune cell states in a tumor-like environment, this work expands our understanding of hemocyte plasticity and establishes *Drosophila* as a genetically accessible model to study the roles of innate immunity in tissue regeneration and tumor progression.

## Results

### Increased abundance and morphological activation of circulating hemocytes in larvae with undead-driven disc overgrowth

We previously reported that an elevated number of plasmatocytes is associated with undead larval eye imaginal discs (genotype: *ey>hid;p35)* (Amcheslavsky et al., 2018; Diwanji and Bergmann, 2020; Fogarty et al., 2016). These disc-associated plasmatocytes exhibit an altered morphology, including cytoplasmic protrusions, in a ROS-dependent manner (Diwanji and Bergmann, 2020; Fogarty et al., 2016). These observations prompted us to also examine hemocytes in circulation in larvae bearing overgrown undead eye discs. Indeed, we found that the total number of circulating hemocytes in larvae with undead discs was significantly higher compared to two control groups, *ey-Gal4* and *ey>p35* (Fig. 1A). Similar findings were also observed in larvae carrying neoplastic tumor discs (*Ras^V12^scrib*) (Pastor-Pareja et al., 2008). Importantly, the increase in circulating hemocyte numbers in larvae with undead discs is dependent on extracellular ROS (eROS) as RNAi knockdown of Duox (the NADPH oxidase that produces eROS in undead discs (Fogarty et al., 2016)) or overexpression of an extracellular catalase (hCatS) significantly reduced this number (Fig. 1A). In contrast, inhibition of downstream JNK signaling through expression of a dominant-negative form (*bsk^DN^*) had no effect on plasmatocyte abundance in *ey>hid;p35* larvae (Fig. 1A).

**Figure 1.**
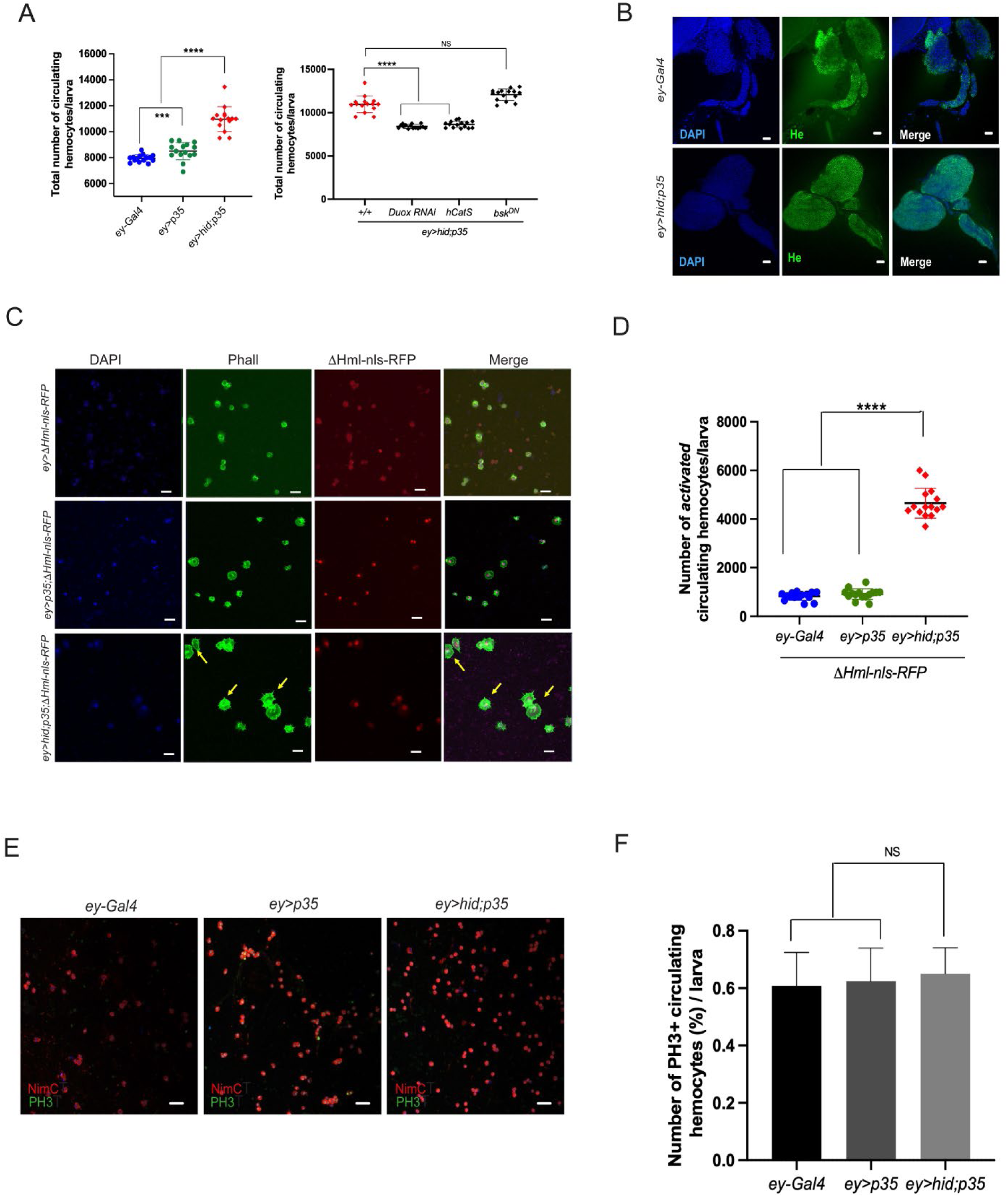
Increased abundance and morphological activation of circulating hemocytes in larvae with undead-driven disc overgrowth. **(A)** Left: Quantification of total circulating hemocytes in third instar larvae co-expressing *hid* and *p35* under control of the *ey-Gal4* driver (*ey>hid;p35*), showing significantly elevated hemocyte numbers compared to control genotypes (*ey-Gal4* and *ey>p35*). Right: RNAi-mediated knockdown of *Duox* or overexpression of an extracellular catalase (*hCatS*) reduces circulating hemocyte numbers, indicating a ROS-dependent effect. Co-expression of a dominant negative JNK transgene (*bsk^DN^*) did not reduce the number of circulating hemocytes. **(B)** Representative confocal images of lymph glands from control (*ey-Gal4*) and *ey>hid;p35* larvae stained with anti-Hemese (He) antibody show intact gland morphology, indicating that the increase in circulating hemocytes is not caused by premature lymph gland rupture. Scale bar: 10µm. **(C)** Representative images of circulating hemocytes labeled with ΔHml-nlsRFP and Phalloidin from larvae of the indicated genotypes reveal morphological alterations in *ey>hid;p35* larvae, characterized by enlarged cell size and extended filopodial protrusions (yellow arrows). Scale bars: 40□μm. **(D)** Quantification of morphologically activated hemocytes per larva reveals that ∼50% of circulating hemocytes in *ey>hid;p35* larvae display this activated morphology. **(E)** PH3 immunostaining of circulating hemocytes to assess mitotic activity. NimC antibody staining was used to identify hemocytes. Scale bar: 5µm. **(F)** Quantification of PH3^+^ hemocytes reveals no significant increase in mitotic index in *ey>hid;p35* larvae compared to controls.

To explain the increased number of hemocytes in larvae with undead discs, we considered that premature rupture of the lymph gland could be responsible as it is known to occur as a response to wasp infection (Banerjee et al., 2019; Bazzi et al., 2018; Leitao et al., 2020; Letourneau et al., 2016). However, lymph gland morphology appears to be intact and shows no signs of disruption in larvae bearing overgrown eye discs (Fig.1B) suggesting that the increased hemocyte count is not directly linked to premature rupture of the lymph gland, but could potentially arise from other mechanisms associated with undead discs. Similar findings were reported in larvae carrying neoplastic *Ras^V12^scrib* tumor discs (Pastor-Pareja et al., 2008).

Circulating hemocytes in *ey>hid;p35* larvae also exhibit a distinct morphology. They are larger and are characterized by long filopodial protrusions (Fig. 1C). We have counted the number of hemocytes with altered morphology per individual larva and found that they represent a large fraction (∼50%) of the total circulating hemocytes (Fig. 1D) indicating a high degree of plasticity of the hemocytes in the presence of undead eye discs. These observations highlight the significant impact of overgrown undead eye discs on the morphology and behavior of circulating hemocytes, suggesting potential functional adaptations in response to these conditions.

Since previous studies have shown that larvae with tumors can produce new hemocytes through *de novo* proliferation (Pastor-Pareja et al., 2008), we investigated whether undead-driven overgrowth triggers similar hemocyte proliferation by PH3 labeling. However, while we detect dividing circulating hemocytes, our analysis revealed no significant increase of mitotic (PH3^+^-labeled) cells within the circulating hemocyte population of larvae with undead discs (Fig. 1E,F). Alternatively, the elevated hemocyte count may result from mobilization of sessile hemocytes into circulation in larvae with undead discs. Such release would indicate that sessile hemocytes contribute to the response to the overgrowth conditions and might represent an adaptive strategy for managing tissue abnormalities.

### Generation of single-cell RNA-seq datasets of hemocytes from larvae carrying undead eye discs

To characterize the systemic immune responses of larvae carrying undead overgrown eye discs (*ey>hid;p35*), we performed microfluidics-based single-cell RNA sequencing (scRNA-seq) using the 10X Chromium platform (Zheng et al., 2017). As controls, we have included the *ey>p35* genotype and *w^1118^* as a wild-type condition in these analyses. For a comprehensive understanding of the complete hemocyte profile in these genotypes, sessile hemocytes were pooled into circulation prior to bleeding (Petraki et al., 2015; Tattikota et al., 2020). 68,354 cells were profiled from 2 replicates per genotype (3 genotypes). We observed a median of ∼17,000 variable transcripts (Suppl Fig S1A) across all genotypes and set the cutoff of cells with mitochondrial RNA up to 20% (Suppl Fig S1B). After this quality control step, we profiled 49,039 cells across the three genotypes by combined integration analyses with Seurat R package using Harmony as batch correcting method (Butler et al., 2018; Korsunsky et al., 2019; Stuart et al., 2019). This approach allowed us to gain insights into the immune response in larvae affected by undead-driven overgrowth.

Initially, dimensional reduction and integration of all cells from the three genotypes identified 20 transcriptionally distinct hemocyte clusters. However, three of these clusters and two others showed highly similar expression profiles (Suppl Fig S1C,D), and were subsequently merged, resulting in a final set of 17 distinct hemocyte clusters (Fig 2A,B). All the cells in these clusters expressed one or more of the pan-hemocyte markers Hml, Srp, NimC1 and He (Suppl Fig 1E,F) (Banerjee et al., 2019), indicating that the cell bleedings used for scRNAseq were largely composed of hemocytes and were not contaminated with other cell types.

**Figure 2.**
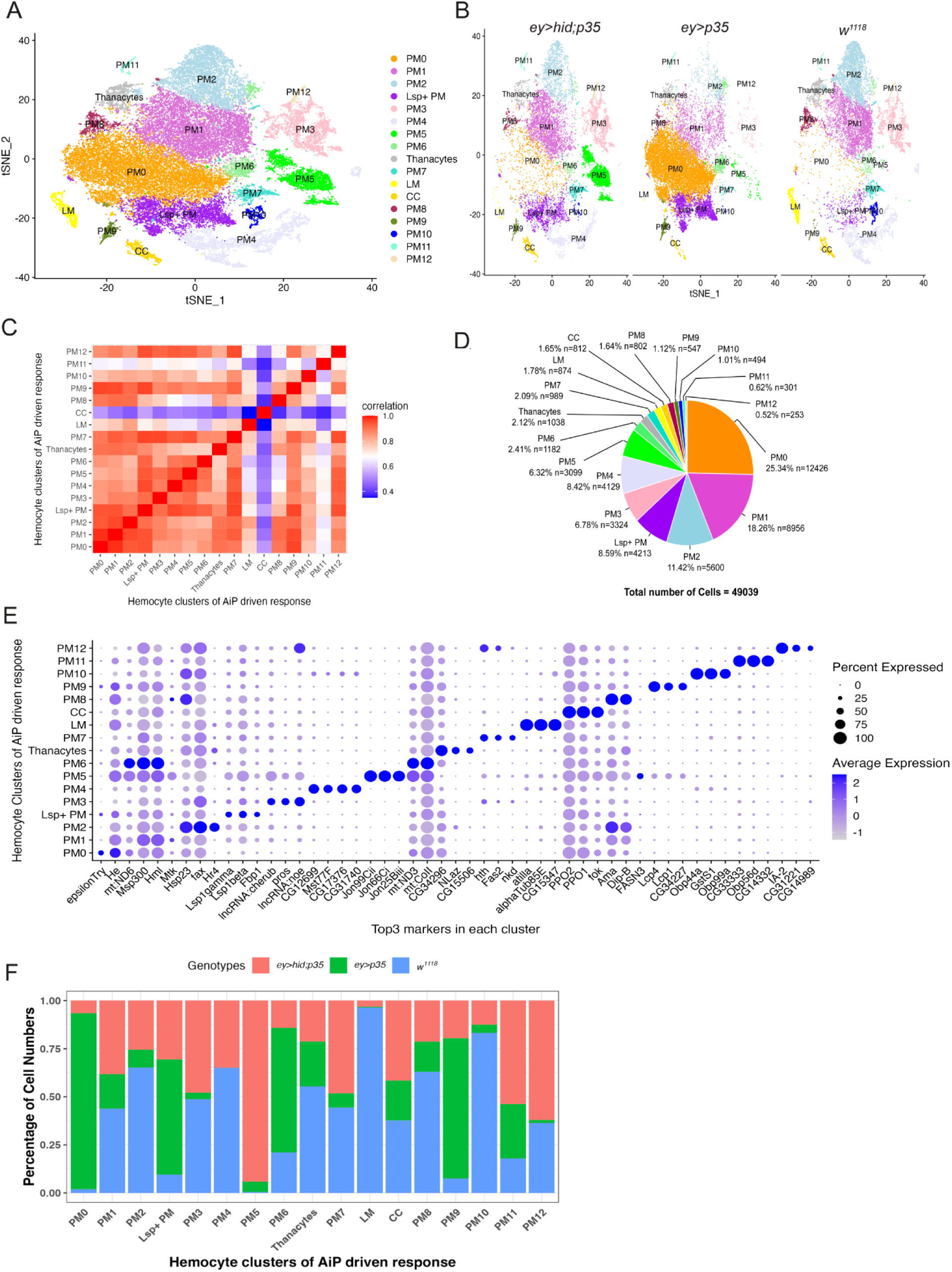
Single-cell RNA sequencing identifies transcriptionally distinct hemocyte populations in larvae with undead-driven overgrowth. **(A)** tSNE projection of 49,039 hemocytes from wild-type (*w^1118^*), *ey>p35*, and *ey>hid;p35* larvae after integration and batch correction using Harmony. Seventeen transcriptionally distinct hemocyte clusters were identified. **(B)** Same tSNE plots as in (A) but showing the distribution of hemocyte clusters separated by genotype. This display illustrates genotype-specific differences in hemocyte composition. **(C)** Heatmap comparing global transcriptional profiles of all clusters. Crystal cells (CCs) and lamellocytes (LMs) display distinct expression profiles compared to other hemocytes. Notably, PM11 also exhibits a unique transcriptional signature, suggesting it represents a functionally distinct plasmatocyte subtype. **(D)** Pie chart indicating relative and total abundance of each hemocyte cluster across the three genotypes. **(E)** Dot plot showing the three highest expressed genes in each cluster. The blue color gradient indicates the expression level of each gene, while the size of each dot reflects the percentage of cells within the cluster expressing the gene. The PM5 cluster is marked by strong expression of Jonah (Jon) family serine proteases. **(F)** Cluster abundance across genotypes. PM5, PM11 and PM12 are enriched in *ey>hid;p35* larvae, while other clusters such as PM0, PM6, and PM10 are under-represented in this condition.

Hemocytes are subdivided into lamellocytes (LM), crystal cells (CC) and plasmatocytes (PM), with the latter making up the majority of hemocytes (Banerjee et al., 2019; Honti et al., 2014; Hultmark and Ando, 2022; Petraki et al., 2015). Consistently, of the 17 hemocyte clusters identified from all combined genotypes, one cluster each represents LM and CC based on marker genes defined by (Tattikota et al., 2020) (Suppl Fig S2C). This classification is further supported by tSNE-presentation using lozenge (lz) and atilla as LM markers, and PPO1 and PPO2 as CC markers (Suppl Fig S2A,B). A heatmap analysis also suggests that CCs and LMs have a very distinct expression profile compared to the other clusters (Fig 2C). There is a strong correlation between the transcriptional profiles of LM and CC in this study compared to previous studies which examined wasp infection in larvae (Cattenoz et al., 2021; Cattenoz et al., 2020; Cho et al., 2020; Leitao et al., 2020; Tattikota et al., 2020) suggesting that these two cell types are not significantly altered by the presence of overgrowing discs in the larvae.

Two additional hemocyte classes, thanocytes and primocytes, have been identified (Fu et al., 2020). We used marker genes defined by Fu et al. to evaluate whether our dataset contained these hemocyte types. While we did not detect primocytes, we identified one cluster that resembles thanocyes by expression of the thanocyte markers *Ance* and *Tep4* (Suppl Fig S2D). We therefore refer to this cluster as thanocytes throughout. The remaining 14 clusters correspond to plasmatocytes (PM).

To classify the plasmatocyte clusters, we examined if they had been previously identified in scRNAseq studies conducted with larval hemocytes (Cattenoz et al., 2020; Cho et al., 2020; Fu et al., 2020; Leitao et al., 2020; Tattikota et al., 2020). Using specific sets of markers identified in these studies, we matched two clusters to previously established populations: Lsp^+^ PM and AMP-PM. The Lsp^+^ PM cluster is distinguished by the expression of larval serum proteins (Lsp) (Suppl Fig S2E) and comprises 8.6 % of the total number of hemocytes (Fig. 2D). Within this cluster, *Fbp1* represents the second most highly expressed gene (Fig 2E) which encodes a receptor for the uptake of Lsp proteins (Burmester et al., 1999).

Enrichment of two to three antimicrobial peptide (AMP)-expressing hemocyte clusters was found to be a characteristic feature during wasp infections (Cattenoz et al., 2020; Tattikota et al., 2020). Enrichment of certain AMPs have also been reported in certain tumor models in Drosophila, although in these studies AMPs were expressed in the fat body, not in hemocytes (Araki et al., 2019; Hanson and Lemaitre, 2020; Parvy et al., 2019). Nonetheless, to explore potential immune activation in our system, we systematically examined AMP expression across all plasmatocyte clusters. In our analysis, we identified a single plasmatocyte cluster (PM5) with prominent expression of the AMPs *Drosomycin* and *Metchnikovin-like* (*Mtkl, CG4236*), along with the immunity factor Relish (Rel) (Suppl Fig S2F). However, although PM5 is the only cluster in our dataset to show induction of AMP genes, we did not classify it as a dedicated AMP-producing population, as it displays a more dominant defining feature discussed below. We did not detect any significant enrichment of CCs, LMs, thanocytes, or Lsp^+^ plasmatocytes within the hemocyte populations that respond to undead disc overgrowth (Fig. 2F).

The remaining 13 plasmatocyte clusters showed distinct transcriptional signatures not described in previous work and were therefore designated as PM0-PM12. This expanded classification reveals the remarkable plasticity and heterogeneity of larval plasmatocytes under undead-driven overgrowth conditions. As key immune system components, hemocytes exhibit transcriptional diversity that reflects their flexible roles in immune surveillance, tissue homeostasis, and possibly tumor suppression or promotion. In the following, we are describing the newly identified plasmatocyte clusters PM0 to PM12. Figure 2E identifies the three highest expressed genes in each cluster.

### Functional annotation of newly identified plasmatocyte clusters obtained from larvae with undead-driven overgrowth

The largest cluster identified was PM0, comprising approximately 25% of all hemocytes analyzed in this study (Fig 2D). Interestingly, PM0 is predominantly present in the control genotype *ey>p35* (>95%) (Fig. 2F), although the underlying reason for this enrichment remains unclear. PM0 plasmatocytes show strong enrichment for gene ontology (GO) terms related to protein processing in the endoplasmic reticulum, cytosolic ribosome, and translation (Suppl Fig S3) indicating high biosynthetic and protein-production capacity. The overrepresentation of PM0 in the control *ey>p35* genotype suggests that its function is unrelated to tissue overgrowth or stress responses, but may instead reflect a general effect of caspase inhibition resulting from *p35* expression. That is intriguing, as *p35* is a protein from baculovirus and thus constitutes a foreign protein within the *Drosophila* proteome. Thus, while *Drosophila* lacks adaptive immunity, these findings raise the possibility that plasmatocytes potentially can detect and transcriptionally respond to viral proteins.

PM1 is the second most abundant plasmatocyte cluster in our dataset, accounting for approximately 18% of all hemocytes (Fig.□2D). Similar to PM0, PM1 is enriched for GO terms cytosolic ribosome, translation, and related protein biosynthesis pathways (Suppl. Fig S3), indicating a high level of basal metabolic activity. In contrast to PM0, which is genotype-specific, PM1 is evenly represented across the three genotypes *w^1118^*, *ey>p35*, and *ey>hid;p35*, suggesting a stable, genotype-independent role in the larval hemocyte pool (Fig 2F).

PM2 comprises a significant portion of the dataset, representing 11.42% of all hemocytes (∼5,600 cells) (Fig.□2D). It is highly enriched in the control genotype *w^1118^*, while substantially underrepresented in larvae with undead disc overgrowth (*ey>hid;p35*) and in *ey>p35* controls (Fig 2F), suggesting that PM2 is preferentially maintained under homeostatic, non-stressed conditions. PM2 is uniquely enriched for GO terms associated with endocytosis, border follicle cell migration, and actin filament binding (Suppl Fig S3). This transcriptional signature points to a functionally dynamic population of plasmatocytes involved in cytoskeletal remodeling and vesicle trafficking. The actin and endocytosis-related terms suggest high motility and a capacity for environmental sensing, surveillance, or clearance of apoptotic debris. Interestingly also, as will be shown in Figure 5, PM2 represents a novel terminally differentiated plasmatocyte population.

PM3 accounts for 6.78% of the total hemocyte population, representing approximately 3,324 cells (Fig 2D). It is evenly distributed between genotype *w^1118^* and *ey>hid;p35*, but strongly underrepresented in *ey>p35* larvae (Fig 2F). PM3 is uniquely enriched for GO terms associated with the precatalytic spliceosome, polytene chromosome, and mRNA binding (Suppl Fig S3). These terms suggest a transcriptionally and post-transcriptionally active population, with enhanced capacity for pre-mRNA splicing, nuclear mRNA regulation, and chromatin-associated processes. The enrichment for polytene chromosome components, typically associated with large-scale transcriptional regulation in *Drosophila*, further implies that PM3 may be engaged in global transcriptional modulation or cell-type-specific gene regulation.

PM4 makes up 8.42% of all cells, or approximately 4,129 individual plasmatocytes (Fig 2D). It is strongly enriched in the *w^1118^* control genotype, where it comprises 68% of its population share, and is significantly depleted in both *ey>hid;p35* (30%) and *ey>p35* (2%) larvae (Fig 2F). This distribution suggests that PM4 may represent a homeostatic or metabolically supportive population that is not maintained under stress or overgrowth conditions. PM4 is enriched for GO terms associated with oxidative phosphorylation, the mitochondrial respiratory chain complex I, and manganese ion binding (Suppl Fig S3). These annotations point toward a heightened mitochondrial activity, likely relying on aerobic respiration for energy generation. The manganese ion binding signature may relate to mitochondrial enzymes such as manganese superoxide dismutase (MnSOD), suggesting a potential role in redox regulation and oxidative stress protection.

PM5 plasmatocytes are significantly enriched in *ey>hid;p35* larvae (Fig 2F) and will be discussed in more detail in the next section.

PM6 constitutes a relatively small but distinct population, comprising 2.41% of all hemocytes, a total of 1,182 cells (Fig 2D). This cluster is most abundant in the *ey>p35* genotype, where it represents 65% of its population, and is present at lower levels in *ey>hid;p35* (15%) and *w^1118^*(20%) larvae (Fig 2F). This distribution suggests that PM6 is specifically enriched in response to expression of the caspase inhibitor p35 and persistent caspase inhibition, but not necessarily dependent on the overgrowth phenotype. PM6 is enriched for GO terms associated with polytene chromosomes, imaginal disc morphogenesis, and dendrite morphogenesis (Suppl Fig S3). These annotations point to a unique transcriptional state potentially linked to developmental signaling, tissue patterning, or cellular remodeling. The enrichment for imaginal disc morphogenesis suggests that PM6 may interact with or respond to tissue-derived cues, while the dendrite morphogenesis signature could reflect cytoskeletal plasticity or signaling integration.

PM7 comprises a modest but clearly defined population, making up 2.09% of all hemocytes, or 989 cells in total (Fig 2D). It is evenly distributed between *w^1118^*and *ey>hid;p35* genotypes (45% each) and relatively underrepresented in *ey>p35* (10%) (Fig 2F). This balanced distribution suggests that PM7 may play a context-independent role under both baseline and tissue stress conditions, but is less favored under caspase inhibition alone. PM7 is enriched for polytene chromosome, nucleoplasm, and negative regulation of DNA-templated transcription (Suppl Fig S3), indicating a distinctive nuclear and transcriptionally repressive state. These features imply that PM7 cells are not actively engaged in high-level biosynthesis, but may instead function in maintaining chromatin structure, silencing gene expression, or stabilizing transcriptional quiescence.

PM8 constitutes a small but distinct plasmatocyte population, representing 1.64% of all hemocytes, or 802 cells (Fig 2D). It is highly enriched in the *w^1118^* control genotype (65%), while underrepresented in both *ey>hid;p35* (20%) and *ey>p35* (15%) backgrounds (Fig 2F). The depletion of PM8 in overgrowth conditions (*ey>hid;p35*) and in the presence of caspase inhibition (*ey>p35*) suggests that this cluster is not maintained under stress, and may instead represent a homeostatic support population. PM8 is transcriptionally enriched for oxidative phosphorylation, cytosolic ribosome, and cytoplasmic translation (Suppl Fig S3), indicating a dual commitment to aerobic energy production and active protein synthesis. These features suggest that PM8 cells are metabolically robust and biosynthetically engaged. The combination of mitochondrial and ribosomal GO terms is reminiscent of PM4, but PM8 exhibits a stronger cytoplasmic translation signature.

PM9 is a small but transcriptionally distinct cluster, comprising 1.12% of total hemocytes, or 547 cells (Fig 2D). It is highly enriched in the *ey>p35* genotype (75%), with lower representation in *ey>hid;p35* (20%) and minimal presence in *w^1118^* (5%) (Fig 2F). This distribution suggests that PM9 arises in response to caspase inhibition, rather than to overgrowth or apoptotic stress per se. PM9 is significantly enriched for GO terms associated with the structural constituent of the chitin-based larval cuticle and chitin-based extracellular matrix (Suppl Fig S3), an unusual profile for hemocytes. Despite this, PM9 cells express canonical hemocyte markers such as Hemolectin (*Hml*) and Hemese (*He*), confirming that they are bona fide plasmatocytes, rather than contaminating cuticle or epithelial cells. The dramatic enrichment of PM9 in *ey>p35* larvae raises the possibility that PM9 represents a stress-induced or mis-differentiated state, potentially triggered by exposure to foreign proteins or disrupted apoptosis. The expression of cuticle-related genes could point to a non-canonical function in extracellular barrier reinforcement, matrix production, or ectopic expression of cuticle-like proteins under conditions of chronic caspase inhibition.

PM10 represents a small plasmatocyte population, comprising just over 1% of the total hemocytes (494 cells) (Fig 2D). It is strongly enriched in the *w^1118^* control genotype (80%), with very limited representation in *ey>p35* (5%) and *ey>hid;p35* (15%) (Fig 2F). This striking distribution suggests that PM10 is a homeostatic plasmatocyte subtype that is preferentially maintained under non-stress conditions and is largely absent in caspase-inhibited or overgrowth contexts. PM10 is enriched for GO terms associated with oxidative phosphorylation, proton-transporting ATP synthase activity, and rotational mechanism (Suppl Fig S3) which are linked to mitochondrial ATP production through the electron transport chain. These terms indicate that PM10 represents a bioenergetically specialized population, likely optimized for efficient aerobic metabolism. The strong enrichment of PM10 in *w^1118^* and its marked depletion in both *ey>p35* and *ey>hid;p35* backgrounds suggest that this cluster is downregulated or outcompeted in the context of stress, apoptosis inhibition, or tissue overgrowth.

PM11 is the second-rarest plasmatocyte cluster, comprising just 0.62% of total hemocytes (301 cells) (Fig 2D). It is preferentially enriched in larvae with undead (*ey>hid;p35*) overgrown discs (55%), with lower representation in *ey>p35* (25%) and *w^1118^* (20%) (Fig 2F). This pattern suggests that PM11 is induced or expanded under chronic apoptotic stress, although not necessarily due to overgrowth conditions. PM11 displays a distinct transcriptional signature (Fig 2C), suggesting that it represents a functionally distinct plasmatocyte subtype. Consistently, GO analysis reveals enrichment for terms such as *response to pheromone* and *chitin-based cuticle development* (Suppl Fig S3). While this signature initially raises the possibility of contamination, the robust expression of the core hemocyte marker Hemolectin (Hml) confirms that PM11 cells are bona fide plasmatocytes (Suppl Fig S1E,F). Its emergence under undead-disc conditions suggests that PM11 is a stress-adapted subpopulation, possibly linked to caspase activity or tissue-derived stress signals. The cuticle-associated gene expression may reflect a developmentally divergent or non-canonical expression program.

PM12 is the least abundant plasmatocyte cluster in the dataset, comprising 0.52% of total hemocytes (253 cells) (Fig 2D). It is enriched in *ey>hid;p35* larvae (60%), moderately present in *w^1118^*(38%), and nearly absent in *ey>p35* (2%) (Fig 2F), suggesting that PM12 emerges in response to persistent apoptotic signaling and overgrowth, but not under caspase inhibition alone. PM12 exhibits an unusual combination of GO terms, including oxidative phosphorylation, neuromuscular junction, and chemical synaptic transmission (Suppl Fig S3). Despite the neuron-associated GO terms, PM12 expresses the hemocyte marker *Hml*, confirming that it is a genuine plasmatocyte population rather than a contaminating neuronal cell type. This raises the intriguing possibility that PM12 represents a functionally divergent hemocyte state, co-opting or mimicking elements of neuronal signaling or synaptic machinery in response to complex apoptotic environments. Its oxidative phosphorylation signature further points to a bioenergetically active state, capable of supporting energy-demanding processes such as vesicular trafficking or secretion. Furthermore, as shown in Figure 5, PM12 does not appear near the terminal ends of any productive lineage trajectories in pseudotime. Instead, it branches early from PM3, the founding progenitor-like state, and enters a developmentally unproductive path, a dead-end trajectory in the pseudotime landscape. This positioning indicates that PM12 is not a transient intermediate, possibly triggered by aberrant or stress-induced differentiation cues. This contrasts with PM5 or PM2, which occupy terminal effector positions.

Finally, the strong enrichment of lamellocytes (LMs) in the *w^1118^*control background (Fig. 2F) is unexpected, as this cell type is typically induced only under infectious or immune-challenging conditions (Banerjee et al., 2019; Koranteng et al., 2022; Monticelli et al., 2024). However, given that LMs constitute only 1.78% of the total hemocyte population in our dataset (Fig. 3D), their elevated numbers in the *w^1118^* background may simply reflect natural variation.

**Figure 3.**
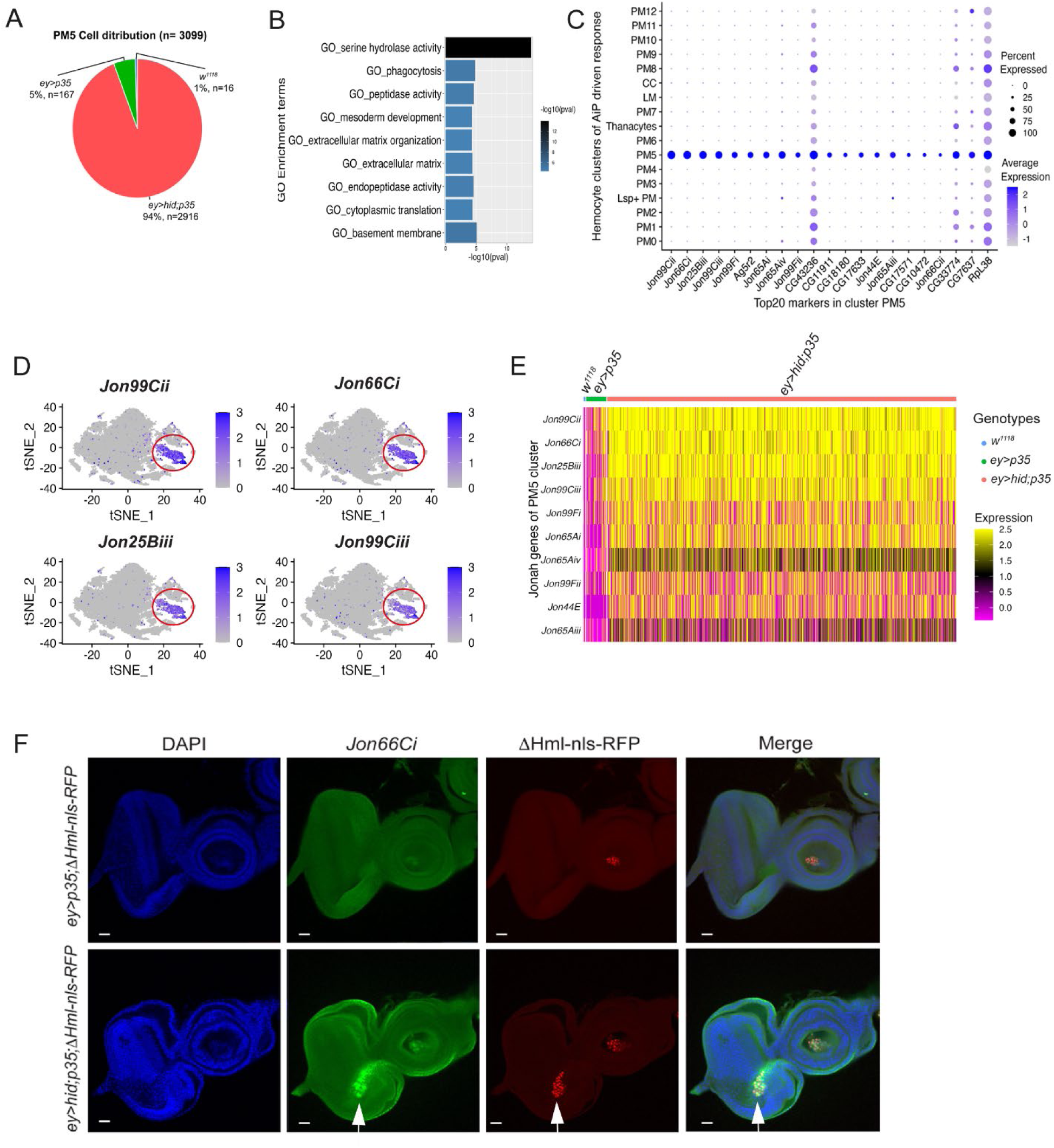
PM5 plasmatocytes are enriched in proteases and respond specifically to undead-driven overgrowth. **(A)** Pie chart showing the strong enrichment of PM5 plasmatocytes in *ey>hid;p35* larvae. **(B)** Gene Ontology enrichment analysis of PM5 marker genes reveals enrichment in serine hydrolase activity. **(C)** Dot plot of the 20 most highly upregulated genes in cluster PM5. Jonah (Jon) proteases dominate this expression profile. The blue color gradient indicates the expression level of each gene, while the size of each dot reflects the percentage of cells within the cluster expressing the gene. **(D)** t-SNE feature plots showing expression patterns of the four most highly expressed *Jonah* family genes (*Jon99Cii*, *Jon66Ci*, *Jon25Biii*, *Jon99Ciii*), specifically restricted to the PM5 cluster (red circle). The blue color gradient indicates the expression level of each gene. **(E)** Heat map depicting the expression levels of the 11 upregulated *Jon* genes in PM5 across all genotypes. **(F)** Representative image showing anti-sense *Jon66Ci*-positive hemocytes (green) localized to undead eye imaginal discs (white arrow). A Δ*Hml-nlsRFP* (red) reporter transgene was used to identify hemocytes. DAPI labeling (blue) marks the outline of the eye-antennal imaginal disc. Genotype of the disc: *ey>hid;p35;* Δ*Hml-nRFP*. Scale bars: 20 µm.

### PM5 plasmatocytes represent a novel subpopulation defined by expression of *Jonah* serine proteases

Of particular interest is cluster PM5 because it is significantly enriched in larvae with overgrown undead eye discs (94%) and thus markedly under-represented in the control genotypes *w^1118^* and *ey>p35* (Fig 2F; 3A). It comprises 3,099 cells representing 6.3% of the total dataset (Fig 2D). GO term analysis of PM5 identifies serine hydrolase and extracellular matrix structural activity as notable features (Fig 3B). Consistently, 16 of the 20 most upregulated genes in PM5 encode proteases (Fig 3C), representing a previously unobserved gene expression profile in plasmatocytes. Remarkably, eleven of these 16 protease-encoding genes belong to the *Jonah* (*Jon*) gene family, which encodes chymotrypsin-like S1 class serine proteases (Fig 2E; Fig 3C-E). The remining five protease-encoding genes include the uncharacterized genes *CG10474*, *CG11911*, *CG17571*, *CG18180* (also encoding S1 serine proteases) and *CG17633* (a metallocarboxypeptidase) (Fig 3D). All these proteases are secreted and function in the extracellular space. The S1 proteases are produced as inactive zymogens and require proteolytic cleavage for activation. The *Jon* proteases are primarily associated with digestive processes in the intestine. However, these proteases may also influence the innate immune response, as their expression levels vary in response to viral, bacterial, fungal and nematode infections (Roxstrom-Lindquist et al., 2004; Sheban et al., 2025; Yadav and Eleftherianos, 2019). Recent work has also detected *Jon* proteases in the larval hemolymph where they act as gelatinases and caseinases (Gatti et al., 2024). These findings support the idea that PM5 plasmatocytes might engage in specialized proteolytic activity.

To better characterize the PM5 cluster, we generated RNA probes targeting *Jon66Ci*, one of the highest expressed *Jon* gene in PM5 hemocytes (Fig 2E; 3C). Because no other PM cluster exhibits a significant induction of *Jon66Ci*, this approach enables the specific identification of PM5 plasmatocytes. Using this marker, *Jon66Ci* probe staining revealed that PM5 plasmatocytes are recruited to undead eye imaginal discs (Fig 3H). This localization suggests that PM5 plasmatocytes may directly contribute to hemocyte-mediated responses at undead imaginal discs.

### PM5 plasmatocytes may facilitate basement membrane repair through coordinated expression of Jonah proteases and matrix components

We previously demonstrated that the basement membrane (BM) of undead imaginal discs sustains damage due to MMP2 proteolytic activity (Diwanji and Bergmann, 2020). Given the localization of PM5 plasmatocytes at undead discs and their enrichment in secreted proteases, particularly Jonah proteases, we hypothesized that PM5 cells contribute to BM repair by combining proteolytic remodeling with the production of new BM components. To explore this possibility, we examined our scRNA-seq dataset for expression of key BM constituents across plasmatocyte clusters, including PM5.

Multiple plasmatocyte clusters exhibited high expression levels of BM components, including Viking (a Collagen IV subunit), Collagen 4α1 (Col4α1), Laminin A (LanA), LanB1, LanB2, and Glutactin (Glt) (Fig. 4A,B). Expression of these genes was particularly elevated in the PM6, PM5 and PM2 clusters, although several other PM clusters also expressed these genes at lower levels (Fig. 4A,B). This transcriptional profile underscores the broader role of plasmatocytes in supplying BM constituents to imaginal discs and other tissues. Among these clusters, only PM5 is strongly enriched in larvae with undead discs, and GO analysis (*extracellular matrix organization)* (Fig. 3B) further highlights a potential role of PM5 in BM remodeling during tissue overgrowth.

**Figure 4.**
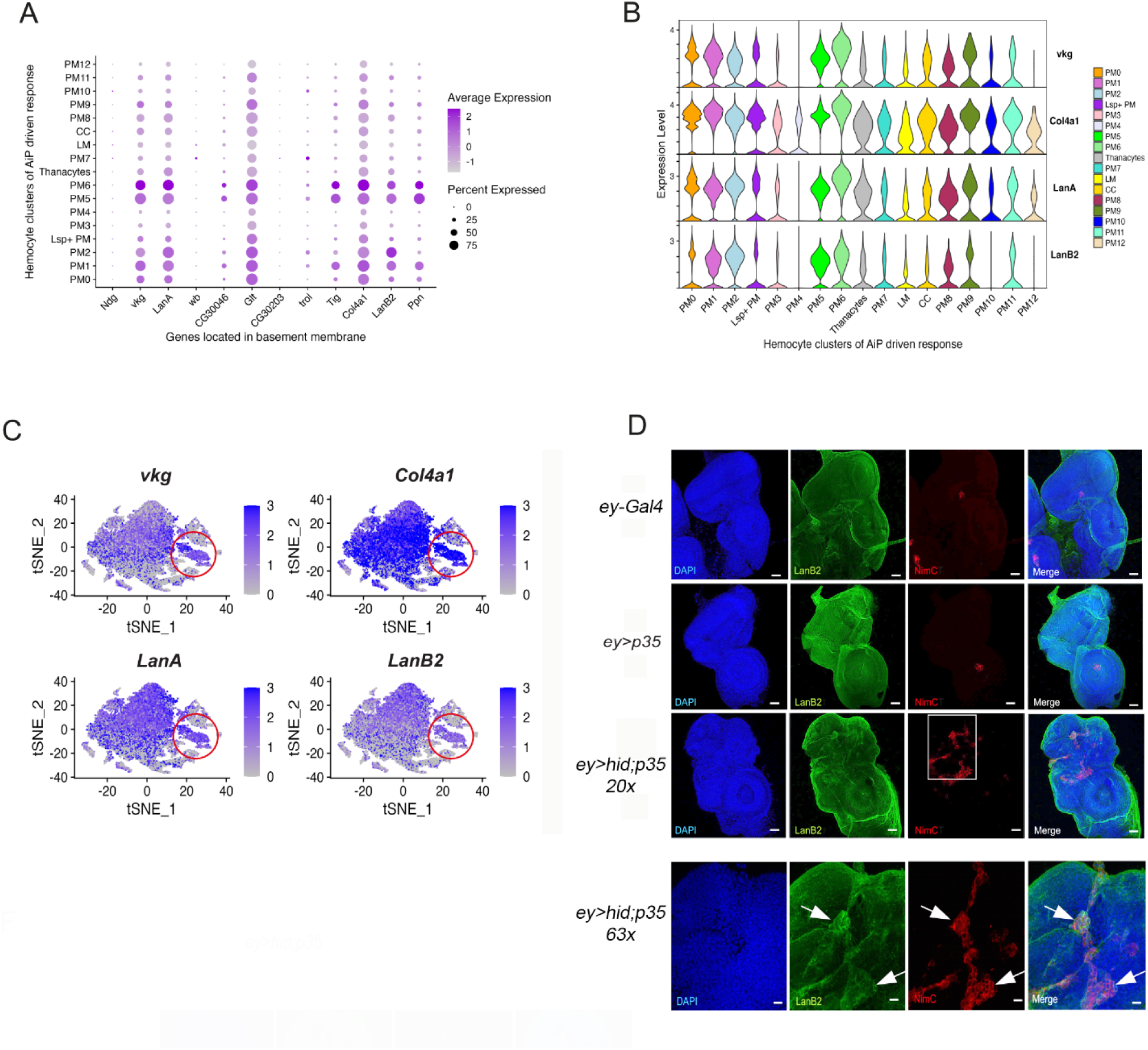
Plasmatocytes may contribute to basement membrane repair through combined expression of Jonah proteases and BM components. **(A)** Dot plot showing expression of key basement membrane (BM) components including *viking* (*vkg*; a Collagen IV subunit), *Col4*α*1*, *LamininA* (*LanA*) *LanB1*, *LanB2*, and *Glutactin* (*Glt*) across plasmatocyte clusters. The purple color gradient indicates the expression level of each gene, while the size of each dot reflects the percentage of cells within the cluster expressing the gene. **(B)** Violin plots showing widespread expression of *vkg*, *Col4*α*1*, *LanA*, and *LanB2* across the plasmatocyte clusters. **(C)** Feature plots visualizing expression of *vkg*, *Col4*α*1*, *LanA,* and *LanB2*, and across the integrated cells, highlighting the broad expression of BM components in plasmatocyte clusters. **(D)** Confocal images of eye-antennal discs from third instar larvae of the indicated genotypes immunostained for LanB2 (green) and the hemocyte marker NimC (red). The white box in the upper *ey>hid;p35* panels is enlarged in the lower panels. LanB2-positive hemocytes are prominently associated with undead eye discs (*ey>hid;p35*) (white arrows), but are absent from discs in the control genotypes. Scale bars upper panels: 20μm; lower panels:100 μm.

Using antibodies raised against LanB2, we examined whether plasmatocytes that express LanB2 are associated with eye imaginal discs. While control genotypes (*w^1118^*and *ey>p35*) did not consistently show LanB2-containing hemocytes associated with eye imaginal discs, we observed robust recruitment of LanB2-producing hemocytes to undead eye imaginal discs (*ey>hid;p35*) (Fig. 4C; see enlargements in Fig. 4D). Given that plasmatocytes provide BM components for functional BM assembly in imaginal discs and other tissues, we conclude that plasmatocytes, in particular PM5 plasmatocytes, are recruited to undead imaginal discs to repair the BM damage characteristic of these discs.

### Pseudotemporal analysis defines plasmatocyte differentiation trajectories

To elucidate the developmental relationships and potential lineage trajectories of hemocyte clusters enriched in response to undead disc overgrowth, we performed pseudotemporal trajectory analysis using Monocle3, a computational framework that reconstructs cell state transitions from static single-cell transcriptomic data (Cao et al., 2019; Trapnell et al., 2014). Pseudotime analysis enables the ordering of individual cells along a putative developmental continuum, allowing the inference of cell fate hierarchies and the identification of both terminally differentiated states and transient intermediate populations.

As the starting point for trajectory inference, we selected cluster PM3, a plasmatocyte population characterized by the highest proportion of cells in the G2/M phase of the cell cycle (Fig 5A). This high mitotic index suggests that PM3 harbors a pool of pluripotent or oligopotent precursor cells, consistent with its elevated expression of *string* (the *Drosophila cdc25* ortholog) (Fig 5B), which promotes entry into mitosis. Based on these features, PM3 was designated as the root node for pseudotemporal ordering.

**Figure 5.**
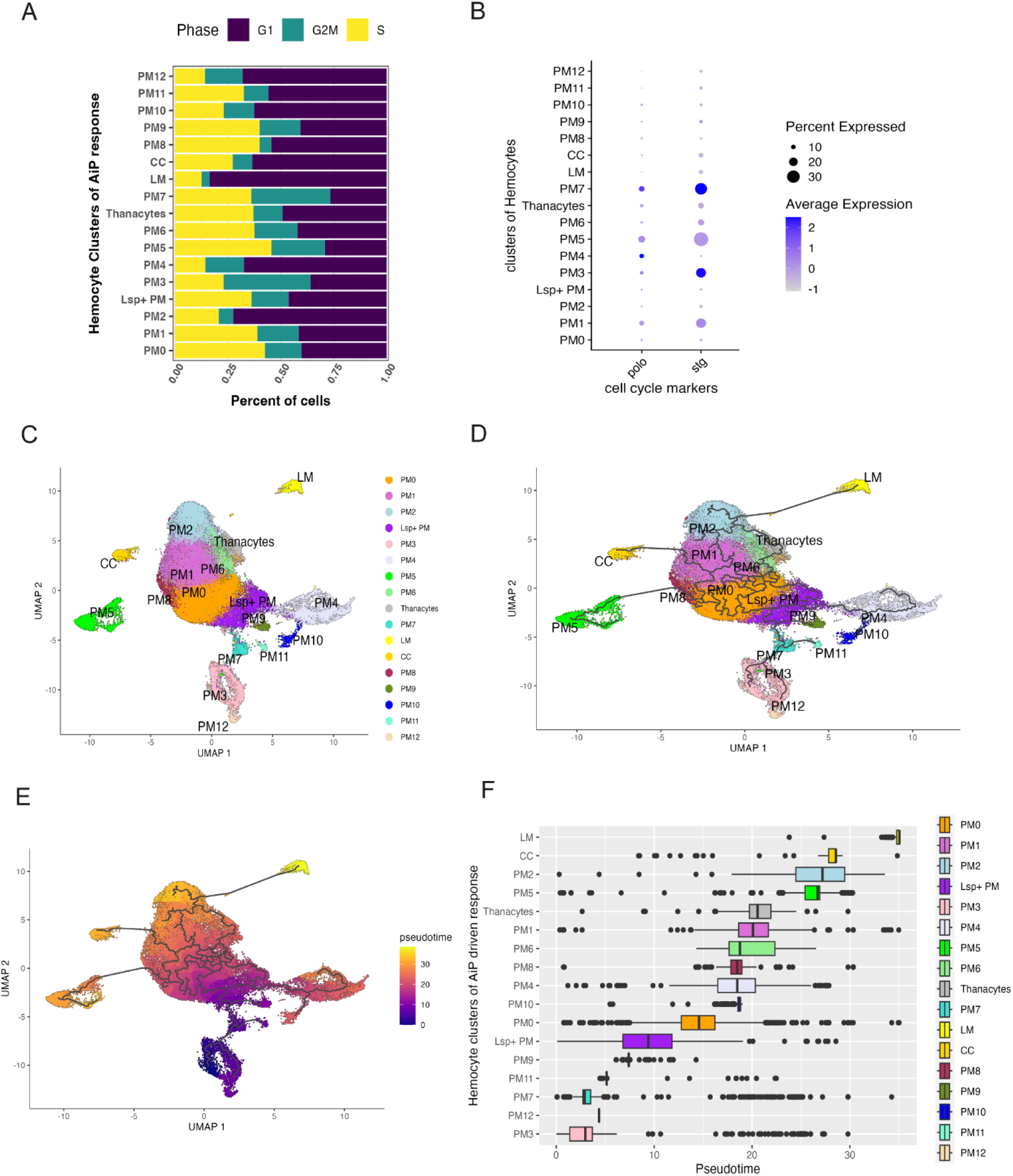
Pseudotemporal analysis defines the developmental plasticity of plasmatocytes and identifies novel terminal states within the hemocyte lineage in response to undead-driven overgrowth. **(A)** Cell cycle phase distribution of each hemocyte cluster, showing the proportion of cells in G1, S, and G2/M phases. PM3 displays the highest proportion of cells in G2/M phase. **(B)** Dot plot showing expression of *polo* (a mitotic regulator) and *string* (the *Drosophila cdc25* ortholog) across the plasmatocyte clusters. The blue color gradient indicates the expression level of each gene, while the size of each dot reflects the percentage of cells within the cluster expressing the gene. **(C)** UMAP visualization of integrated hemocyte clusters. **(D)** Developmental trajectories of each individual cluster inferred using Monocle 3, rooted in the PM3 cluster. **(E)** Pseudotime map generated using Monocle3 reveals crystal cells (CCs) and lamellocytes (LMs) at distinct terminal branches. PM2 and PM5 lie at the terminus of unique branches, suggesting previously uncharacterized terminal differentiation states within the plasmatocyte lineage. **(F)** Linear hierarchical pseudotime representation of hemocyte clusters positioned along the inferred developmental trajectory. Colored boxes indicate the identity and relative pseudotime placement of each cluster. PM3, designated as the root, appears at the earliest point in pseudotime and likely represents an immature progenitor state. Crystal cells (CCs), lamellocytes (LMs), PM2 and PM5 occupy terminal positions, consistent with their status as differentiated end states.

The pseudotime trajectory revealed that crystal cells and lamellocytes occupy distinct terminal branches, consistent with their roles as specialized, terminally differentiated hemocyte lineages (Fig 5C-F). This observation is in agreement with previous scRNAseq analyses of *Drosophila* hemocytes under immune challenge and developmental differentiation conditions (Cattenoz et al., 2020; Cho et al., 2020; Coates et al., 2021; Fu et al., 2020; Girard et al., 2021; Leitao et al., 2020; Tattikota et al., 2020).

In addition to these classical lineages, our analysis of plasmatocyte clusters obtained from larvae with undead disc overgrowth revealed that PM2 and PM5 also lie at the end of distinct pseudotemporal branches, suggesting that they represent the most differentiated plasmatocyte subpopulations (Fig 5C-F). Unlike previously described terminal states, PM2 and PM5 have not been functionally characterized as such, and their position at the trajectory endpoint identifies them as a previously unrecognized terminal differentiation state within the plasmatocyte lineage, particularly in the context of AiP-driven undead disc overgrowth.

Cluster PM5 is of particular interest as it is strongly enriched in larvae with undead overgrown eye discs and defined by high expression of Jonah proteases (Fig.□3) as well as genes associated with basement membrane remodeling (Fig.□4). The terminal positioning in pseudotime suggests that PM5 cells have reached a mature state tailored for damage response and matrix repair in response to chronic overgrowth and stress.

In addition, PM2 represents a second terminally differentiated plasmatocyte population that resides at the endpoint of the pseudotime trajectory (Fig.□5E, F). Both clusters express basement membrane (BM) components at elevated levels (Fig. 4), suggesting a shared capacity for extracellular matrix association or structural support. However, PM2 differs markedly from PM5 in both functional annotation and contextual enrichment. Whereas PM5 is strongly induced under AiP-driven overgrowth and expresses high levels of proteolytic genes such as Jonah family serine proteases, PM2 lacks protease expression and is instead enriched in the *w^1118^* control genotype (70%), but underrepresented in both *ey>hid;p35* (25%) and *ey>p35* (5%) backgrounds. Gene ontology analysis of PM2 reveals terms including *endocytosis*, *border cell migration*, and *actin filament binding* (Suppl Fig S3) suggesting a role in cytoskeletal remodeling and potentially in matrix surveillance or cell motility. We propose that PM2 may contribute to tissue homeostasis through endocytic clearance or structural maintenance, operating as a specialized, non-inflammatory effector population under steady-state conditions. Its marked depletion in overgrowth-associated genotypes supports the idea that this lineage is either suppressed or redirected in response to chronic tissue stress.

## Discussion

In this study, we investigated the diversity and differentiation trajectories of hemocytes in *Drosophila* larvae bearing undead overgrown eye imaginal discs, a model of apoptosis-induced proliferation (AiP). AiP is a compensatory mechanism in which dying cells release mitogens to promote their own replacement (Bergmann, 2025; Bergmann and Fan, 2025; Diwanji and Bergmann, 2019; Fogarty and Bergmann, 2017). When these apoptotic cells are rendered immortal or “undead,” this regenerative program becomes dysregulated and can drive tissue overgrowth. This overgrowth phenotype of undead tissue has proven to be a powerful experimental model for uncovering the cellular and molecular mechanisms underlying AiP. Previous studies showed that hemocytes, in particular plasmatocytes, are recruited to undead tissue in a ROS-dependent manner and exhibit altered morphology suggestive of activation (Diwanji and Bergmann, 2020; Fogarty et al., 2016). The degree of hemocyte activation varies between different tumor and overgrowth models of eye-antennal imaginal discs. Neoplastic models, such as *Ras^V12^scrib* and *vps25* mosaic discs, are associated with hemocytes exhibiting a highly activated morphology, whereas hyperplastic models, including *hippo* mosaics and *eyeful* discs, show only minimal hemocyte activation (Diwanji and Bergmann, 2020; Perez et al., 2017). In this regard, the hemocyte response to undead-driven overgrowth resembles more closely that seen in neoplastic models, suggesting that undead tissue elicits a similarly robust immune activation state compared to neoplastic tissue. Here, we extended these observations through transcriptome-wide single-cell RNA sequencing (scRNA-seq) of circulating and sessile hemocytes obtained from larvae with undead overgrown discs, and uncovered a broad repertoire of hemocyte subtypes, novel effector states, and unexpected differentiation trajectories.

We showed that larvae bearing undead overgrown discs exhibit a significant increase in circulating hemocytes, accompanied by striking morphological changes, including enlarged cell size and extended filopodia in ∼50% of the hemocyte population (Fig 1). The expansion and morphological activation of circulating hemocytes in larvae with undead discs are reminiscent of inflammatory mobilization seen in tumor-bearing vertebrates (Kloosterman and Akkari, 2023; Ma et al., 2022; van Vlerken-Ysla et al., 2023). However, we observed no evidence for uncontrolled proliferation of hemocytes, nor for lymph gland rupture (Fig 1), indicating that the observed hemocyte expansion potentially results from a regulated redistribution of sessile immune cells, rather than a generalized stress response. This increase is dependent on extracellular ROS and damage of the basement membrane (Diwanji and Bergmann, 2017; Diwanji and Bergmann, 2018; Diwanji and Bergmann, 2020; Fogarty et al., 2016), and mirrors immune responses observed in neoplastic tumor models (e.g., *Ras^V12^scrib*) (Hirooka et al., 2025; Pastor-Pareja et al., 2008; Perez et al., 2017), suggesting that undead discs elicit a tumor-like systemic immune response.

Our scRNA-seq analysis provides a comprehensive map of hemocyte heterogeneity in *Drosophila* larvae containing undead imaginal discs. In total, we identified 17 distinct hemocyte clusters, including previously described crystal cells (CCs), lamellocytes (LMs), thanocytes and Lsp^+^-expressing plasmatocytes (Fig 2). More strikingly, our analysis reveals an unexpected degree of transcriptional and functional heterogeneity, developmental states and genotype-specific enrichment patterns within the plasmatocyte lineage, uncovering 13 distinct previously undescribed PM clusters (PM0-PM12).

Several PM clusters (e.g., PM0, PM1, PM4, PM8, PM10) were enriched under control conditions and exhibited transcriptional signatures related to translation, oxidative phosphorylation, and general matrix production, consistent with homeostatic or supportive roles in healthy larvae. Several rare or stress-associated clusters, including PM6, PM9, PM11, and PM12, exhibit transcriptional programs not typically associated with hemocytes, such as morphogenetic, cuticle, or even neuron-like gene signatures, yet clearly express core plasmatocyte markers like *Hml*, suggesting either adaptive or mis-directed differentiation under persistent apoptotic or caspase-inhibited conditions. Notably, PM11 and PM12 do not lie at terminal positions in the pseudotime landscape. PM12, in particular, branches into a non-productive, dead-end trajectory downstream of PM3, reflecting a potentially mal-adaptive response to chronic apoptotic signaling. These findings underscore the remarkable plasticity of the plasmatocyte lineage and reveal how apoptotic signaling and caspase inhibition dynamically reshape the landscape of hemocyte differentiation and function *in vivo* that go far beyond phagocytosis or antimicrobial defense. This expanded classification reveals substantial plasmatocyte heterogeneity and underscores the transcriptional plasticity of the hemocyte response under conditions of apoptosis-induced tissue overgrowth.

To explore hemocyte differentiation trajectories, we applied pseudotime analysis using Monocle 3, with PM3 selected as the root node based on its high G2/M index and elevated expression of *string* (the *cdc25* ortholog), suggesting a mitotically primed, oligopotent progenitor-like identity. The pseudotime trajectory revealed that lamellocytes and crystal cells reside at terminal endpoints, consistent with their status as differentiated immune effectors, and corroborating earlier scRNA-seq findings from wasp infection models (Cattenoz et al., 2020; Tattikota et al., 2020). Importantly, PM2 and PM5 emerge as previously unrecognized terminally differentiated plasmatocyte subtypes at the end of separate pseudotemporal branches, defined by a unique transcriptional signature.

Of particular interest, PM5 emerged as a terminally differentiated effector population that is selectively enriched in larvae with undead disc overgrowth. PM5 plasmatocytes display a unique transcriptional profile characterized by high expression of Jonah (Jon) family serine proteases, as well as basement membrane (BM) components such as Collagen IV (Viking), Laminins, and Glutactin. The Jon gene family consists of 16 members distributed in seven genomic clusters across multiple chromosomes, encoding proteases of ∼28 kDa from diverse subfamilies (Carlson and Hogness, 1985). These proteases exhibit dynamic expression patterns throughout *Drosophila* development, with broad activity during larval stages that diminishes by the end of the third instar. In adults, expression resumes in tissues such as the midgut, fat body, imaginal discs, and salivary glands (Carlson and Hogness, 1985). They are traditionally linked to digestive processes, but have also been shown to respond transcriptionally to infections caused by viruses, bacteria, fungi, and nematodes (Roxstrom-Lindquist et al., 2004; Sheban et al., 2025; Yadav and Eleftherianos, 2019). Moreover, recent work has detected Jon proteases in the larval hemolymph, where they function as gelatinases and caseinases (Gatti et al., 2024).

These observations suggest that in addition to their digestive roles, Jon proteases may contribute to extracellular matrix (ECM) remodeling under conditions of stress or tissue overgrowth. In line with this model, *in situ* hybridization analysis revealed that *Jon66Ci*-expressing PM5 plasmatocytes are recruited to undead eye discs, placing them directly at sites of overgrowth and tissue (BM) injury. This spatial localization, combined with their strong expression of both proteases and BM components supports a dual role for PM5 in ECM turnover and BM repair. Together, these findings indicate that hemocytes can dynamically transition into tissue-repairing states, in which they both supply BM components and remodel the ECM through localized proteolysis in response to persistent apoptotic signaling.

PM5 also stands out as the only PM cluster expressing antimicrobial peptides (AMPs) such as Drosomycin and Metchnikovin-like (Mtkl), along with the immune regulator Relish, indicating an integrated immune-effector phenotype. Therefore, rather than forming a dedicated AMP cluster, immune activation in undead larvae appears to be incorporated into the multifunctional PM5 identity. We propose that PM5 represents a flexible, damage-responsive arm of the plasmatocyte lineage, capable of mounting a coordinated response to persistent tissue stress by engaging both immune and repair pathways. Analogous to M2-like tumor-associated macrophages (TAMs) in mammals, PM5 may promote localized ECM remodeling and immune modulation in response to persistent caspase signaling. The behavior of PM5 cells illustrates how innate immune cells can modulate their environment in response to pathological growth. These findings expand our understanding of plasmatocyte plasticity and highlight the existence of specialized hemocyte states beyond classical phagocytosis or antimicrobial defense.

In contrast to PM5, the PM2 plasmatocytes define a second novel terminally differentiated lineage that appears to support homeostatic, rather than damage-induced, functions. PM2 cells constitute over 11% of the total hemocyte population and are highly enriched in the *w^1118^* control genotype (70%), but underrepresented in larvae with undead discs (*ey>hid;p35*, 25%) and even more so in *ey>p35* controls (5%). Like PM5, PM2 expresses multiple BM components at high levels, suggesting a shared capacity for extracellular matrix engagement. However, PM2 lacks expression of *Jon* proteases and other tissue-remodeling enzymes, and does not express AMPs or immune regulators. Instead, gene ontology analysis highlights functions related to *endocytosis*, *border cell migration*, and *actin filament binding*, implicating PM2 in cytoskeletal dynamics, matrix surveillance, or clearance functions. We propose that PM2 represents a homeostatic effector plasmatocyte lineage that contributes to tissue maintenance under steady-state conditions. Its reduced abundance under overgrowth or caspase-inhibited conditions suggests that chronic stress either suppresses its differentiation or redirects progenitor cells toward alternative, stress-adapted fates such as PM5.

Another key insight is that not all classical immune cell types contribute equally to the response. Lamellocytes and crystal cells, while present, are not enriched under undead growth conditions, and their transcriptional profiles remained largely unchanged. This stands in contrast to infection models where lamellocyte expansion is robust (Cattenoz et al., 2020; Cho et al., 2020; Coates et al., 2021; Girard et al., 2021; Leitao et al., 2020; Tattikota et al., 2020). Furthermore, as stated above, we did not identify any specialized AMP-producing clusters in larvae with undead overgrowth. These findings suggest that the immune landscape triggered by undead overgrowth appears qualitatively distinct from canonical inflammatory responses. Rather than mobilizing traditional immune effector types, the response favors specialized subsets of plasmatocytes, such as PM5, that integrate both matrix repair and innate immune functions into a context-specific effector program.

In conclusion, our study presents the first comprehensive single-cell atlas of hemocytes in the context of AiP-induced tissue overgrowth and uncovers striking hemocyte plasticity under non-infectious stress. We reveal a highly dynamic response characterized by increased hemocyte numbers, activation of specialized plasmatocyte subtypes, and the emergence of novel, terminally differentiated effector states such as PM5. These findings broaden our understanding of the *Drosophila* immune system by demonstrating that hemocytes, beyond immune surveillance, play active roles in matrix remodeling and repair, functioning as both structural and regulatory contributors to tissue homeostasis.

The discovery of extensive plasmatocyte heterogeneity and unexpected differentiation trajectories in response to tissue overgrowth opens several promising avenues for future investigation. One key priority will be to functionally validate the specialized roles of newly identified clusters, particularly terminally differentiated populations like PM5, in basement membrane remodeling, immune activation, and the regulation of AiP. Similarly, the molecular cues that govern trajectory branching from mitotically active progenitors like PM3 into effector (e.g., PM5) versus unproductive fates (PM12) remain unknown. Dissecting the upstream signals, whether local, systemic, or damage-derived, that drive these lineage decisions will be essential to understanding how innate immune cells interpret and respond to chronic stress. In parallel, exploring whether similar transcriptional states arise in other models of regeneration, infection, or tumorigenesis will help define conserved versus context-specific features of hemocyte plasticity. Altogether, this work lays the foundation for mechanistic studies that link hemocyte transcriptional states to function, and for leveraging *Drosophila* as a powerful genetic model to uncover fundamental principles of immune-tissue crosstalk in development, regeneration, and disease.

## Material and Methods

### Fly stocks and reagents

Third instar *Drosophila melanogaster* larvae of the genetic backgrounds *w^1118^*, *ey>p35* (complete genotype: *ey-Gal4 UAS-p35*), and *ey>hid, p35* (*UAS-hid; ey-Gal4 UAS-p35*) were used for the preparation of single hemocytes. The transgene Δ*Hml-nRFP* was used to identify hemocytes associated with imaginal discs. *UAS-Duox^RNAi^*, *UAS-hCatS* (both a kind gift of Dr. Won-Jae Lee, (Ha et al., 2009) and UAS-*bsk^DN^ (BL6409)* were used for hemocyte counts. All stocks and crosses were maintained on standard fly food at 25°C.

### Hemocyte preparation, sequencing and data processing

For each biological replicate (n = 2 per genotype), hemocytes were isolated from approximately 50-70 third instar *Drosophila* larvae, following the protocols described by (Tattikota et al., 2020; Tattikota and Perrimon, 2021). Single-cell libraries were prepared using 10x Genomics Single Cell 3’ v3.1 chemistry, following the manufacturer’s instructions. Indexed cDNA libraries from six different samples were pooled and sequenced on a single lane of an Illumina NovaSeq S4 flow cell, using paired-end sequencing (2 × 150 base pairs).

### Cell clustering and data integration

Raw sequencing data were processed using the Cell Ranger software (v6.0.1, 10x Genomics) and aligned to the *Drosophila melanogaster* reference genome (BDGP6_ens95) via the DolphinNext platform (Yukselen et al., 2020) at the University of Massachusetts Medical School. Across all six samples -comprising two biological replicates per genotype -68,354 cells were initially identified. Downstream analysis and clustering were carried out using the R-based Seurat package (v4.1; RRID:SCR_016341) (Butler et al., 2018; Stuart et al., 2019). Quality control filtering retained cells with expression of at least 200 genes and no more than 20% mitochondrial gene content, yielding a final dataset of 49,039 high-quality cells. Each dataset was independently log-normalized, and ∼2,000 variable genes were identified using the “vst” (variance stabilizing transformation) method.

For integrated analysis across all conditions, filtered datasets were merged based on shared genes into a single Seurat object and integrated using Harmony (Korsunsky et al., 2019) to correct for batch effects. Clustering was performed using the first 20 principal components at a resolution of 0.5. Additional clustering was performed separately for control and overgrowth (undead) samples to assess condition-specific differences.

For visualization, the clustered cells were embedded in two-dimensional space using t-distributed Stochastic Neighbor Embedding (t-SNE) with default parameters. Dot plots and heatmaps were generated using Seurat’s DotPlot and DoHeatmap functions, respectively, with heatmaps stratified by condition.

### Pseudotemporal ordering of cells using Monocle 3

Pseudotime analysis was performed using Monocle 3 (Cao et al., 2019; Trapnell et al., 2014) (https://github.com/cole-trapnell-lab/monocle3) to reconstruct cell differentiation trajectories from the integrated overgrowth dataset. The Seurat object containing the integrated UMAP-clustered data was converted to a Monocle 3 CellDataSet (cds) object using the SeuratWrappers R package, which facilitates interoperability between Seurat and Monocle. Cell trajectories were learned using Monocle’s align_cds and learn_graph functions. Cluster PM3 was selected as the root (starting point) for pseudotime alignment, based on its high average expression of the mitotic genes *polo* and *stg* (Leitao et al., 2020; Tattikota et al., 2020) and high G/M score, consistent with a highly proliferative, early progenitor-like state. Gene expression dynamics along pseudotime were extracted using the plot_genes_in_pseudotime function and visualized using ggplot2 (v3.2.1).

### Data files

The original FASTQ files and UMI-filtered gene expression count matrices in MatrixMarket Exchange (MTX) format [see https://math.nist.gov/MatrixMarket/info.html for details] will be deposited in the NCBI Gene Expression Omnibus (GEO).

### Gene set enrichment analysis

Gene set enrichment analysis (GSEA) was performed on cluster-specific marker genes showing positive fold change using the easyGSEA site (https://tau.cmmt.ubc.ca/eVITTA/easyGSEA/) (Cheng et al., 2021). The analysis included curated gene sets from Gene Ontology (GO), Biological Process (BP), Molecular Function (MF), and Cellular Component (CC) categories, as well as pathway annotations from KEGG and Reactome databases. Enrichment significance was calculated based on the p-value, and the strength of enrichment was visualized in bar plots as the negative log□□ of the p-value (–log□□□ p) using the ggplot2 (v3.2.1) function of R package.

### Dissection and immunostaining of larval lymph glands

Third instar larval lymph glands were dissected in ice cold 1xPBS buffer and fixed in 4% paraformaldehyde (PFA) for 40 minutes. Following fixation, lymph glands were permeabilized, blocked and incubated with anti-Hemese (He) antibody at 1:200 dilution. After washing, samples were stained with secondary antibody, mounted, and imaged by confocal microscopy.

### Immunostaining of circulating hemocytes

Circulating hemocytes were collected from third instar larvae after brief vortexing them to dislodge sessile hemocytes. Individual larvae were bled into 0.2□mL of 1× PBS,. The resulting cell suspension was transferred into Lab-Tek II chambered coverglass wells (VWR, Cat# 62407-056). The resulting cell suspension was allowed to settle for 30 minutes. Hemocytes were fixed in 4% paraformaldehyde for 20 minutes. Cells were permeabilized with 0.1% PBST (1× PBS containing 0.1% Triton X-100) for 10 minutes and subsequently blocked with 5% BSA in PBST for 20 minutes. Primary antibody incubations were performed overnight at 4□°C using the following dilutions: anti-Hemese (1:200) and anti-phospho-Histone H3 (PH3, 1:2,000). The next day, samples were incubated with appropriate secondary antibodies (1:500 dilution) for 2 hours at room temperature. Phalloidin-GFP (1:400 dilution) was included to label F-actin structures. After three washes with PBST, mounting medium containing DAPI was added directly to the wells. Confocal imaging was performed on a Zeiss LSM700 microscope. The hemocyte numbers were calculated from the captured images with imageJ.

### Immunostaining of eye imaginal discs

Immunostaining was performed on fixed third instar larval eye-antennal discs following standard protocols as described in (Fogarty and Bergmann, 2014). Discs were incubated with anti-NimC antibody (1:300) (Kurucz et al., 2007), and anti-Laminin B2 (LanB2) (Abcam, # ab47651) at a 1:600 dilution. Fluorescently conjugated secondary antibodies were purchased from ThermoFisher and used according to the manufacturer’s recommendations.

### Fluorescence *in situ* Hybridization

Fluorescent anti-sense and sense RNA probes targeting *Jon66Ci* were generated using the Molecular Probes™ FISH Tag™ RNA Multicolor Kit with Alexa Fluor™ dyes (ThermoFisher, Cat# F32956), following the manufacturer’s instructions. Probes was prepared from DGRC clone BS13309 (DGRC Stock 1632030 ; https://dgrc.bio.indiana.edu//stock/1632030; RRID:DGRC_1632030) with the primers, FW: CCAAGTTGTTTCTGCAACATGA and RV: TCGTTGTAGTCGTTGTAGCGATC. The sequence obtained was cloned into pGEM®-T Easy Vector (Promega, A1630) and the sequence was verified. The verified sequences were labeled as per the manufacturer’s instructions. Third instar larval eye imaginal discs were hybridized overnight at 50□°C in hybridization buffer. The following day, samples underwent a series of washes in decreasing concentrations of SSC (Sodium Saline Citrate) buffer. Discs were then mounted and imaged according to the recommended protocol provided by the manufacturer.

### Statistics

All statistical analyses including quantification of fluorescence intensity and cell counts were conducted using GraphPad Prism 9 software. Error bars represent either the standard error of the mean (SEM) or standard deviation (SD), as indicated in the corresponding figure legends. For comparisons between two groups, unpaired two-tailed Student’s *t*-tests were used. When comparing more than two groups, one-way ANOVA was applied. Statistical significance is indicated as follows: p□<□0.05 (*), p□<□0.01 (**), p□<□0.001 (***), and p□<□0.0001 (****).

## Acknowledgements

We thank our colleague Dr. Nathan Lawson and his team for granting us access to the 10x Genomics platform, which enabled the single-cell RNA-seq experiments described in this study. We are grateful to Dr. István Andó for providing the anti-He and anti-Hml antibodies and to Dr. Won-Jae Lee for providing *UAS-Duox^RNAi^* and *UAS-hCatS*. We also thank Dr. Onur Yukselen, Dr. Alper Kucukural, Dr. Kai Hu and Dr. Julie Zhu for their assistance with bioinformatics analysis. Fly stocks were obtained from the Bloomington *Drosophila* Stock Center (NIH P40OD018537). *Drosophila* genomic clones were obtained from Drosophila Genomics Resource Center (NIH Grant 2P40OD010949). This work was supported by the National Institute of General Medical Sciences (NIGMS) of the National Institutes of Health (NIH) under award number R35GM118330. The content is solely the responsibility of the authors and does not necessarily represent the official views of the funders.

## Author contributions

PY: Conceptualization, Data curation, Formal Analysis, Investigation, Methodology, Validation, Writing – original draft.

AB: Conceptualization, Data curation, Formal Analysis, Funding acquisition, Resources, Supervision, Writing -review and editing.

