## Supplemental Figure S1-S3 for "Single-cell RNA analysis reveals unexpected hemocyte plasticity and immune cell specialization in a *Drosophila* overgrowth model"

Fig S1: Quality control and Specicity of Hemocyte preparation for Sc-seq data analysis

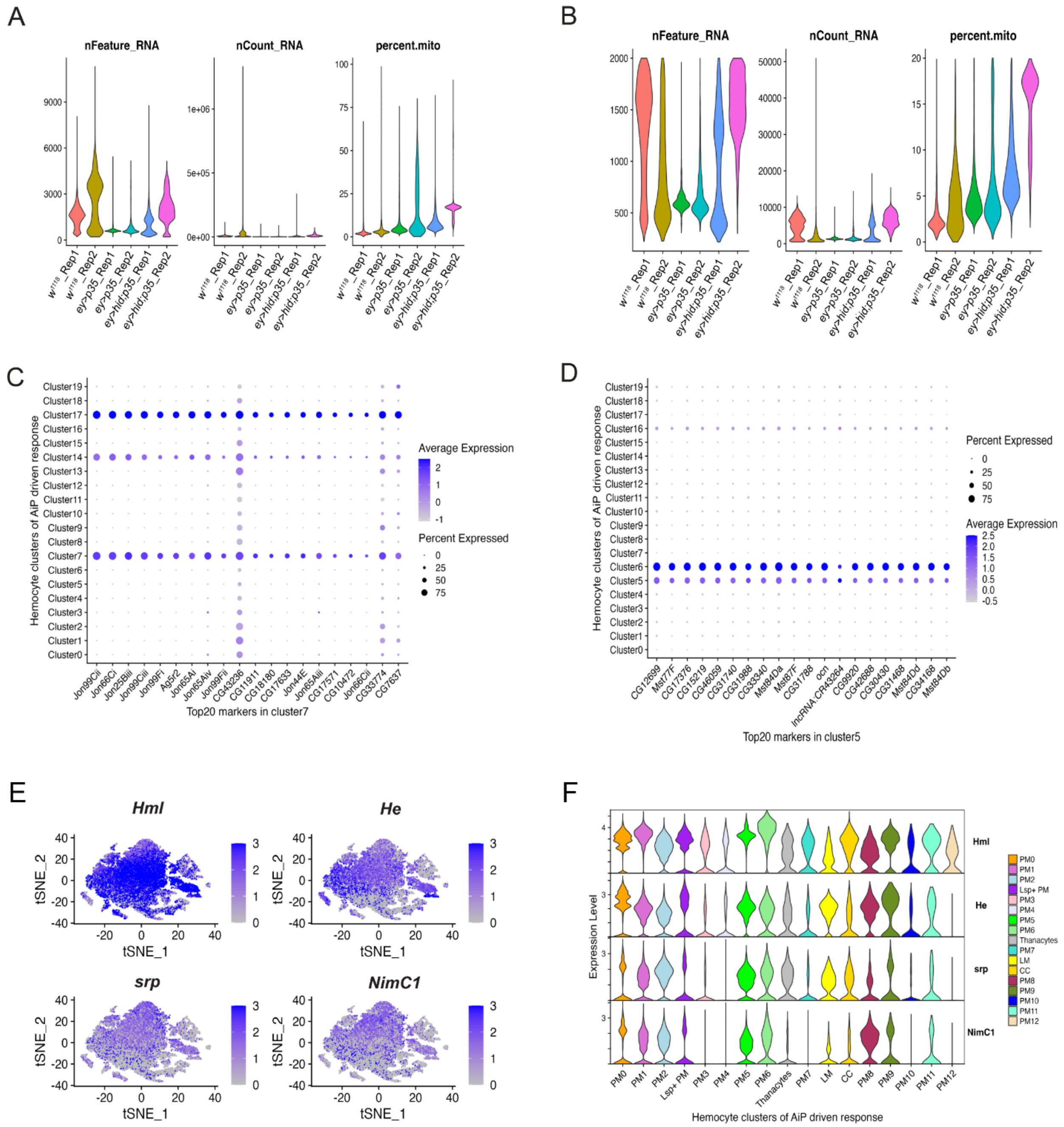

Fig S2: Identification of known hemocyte types

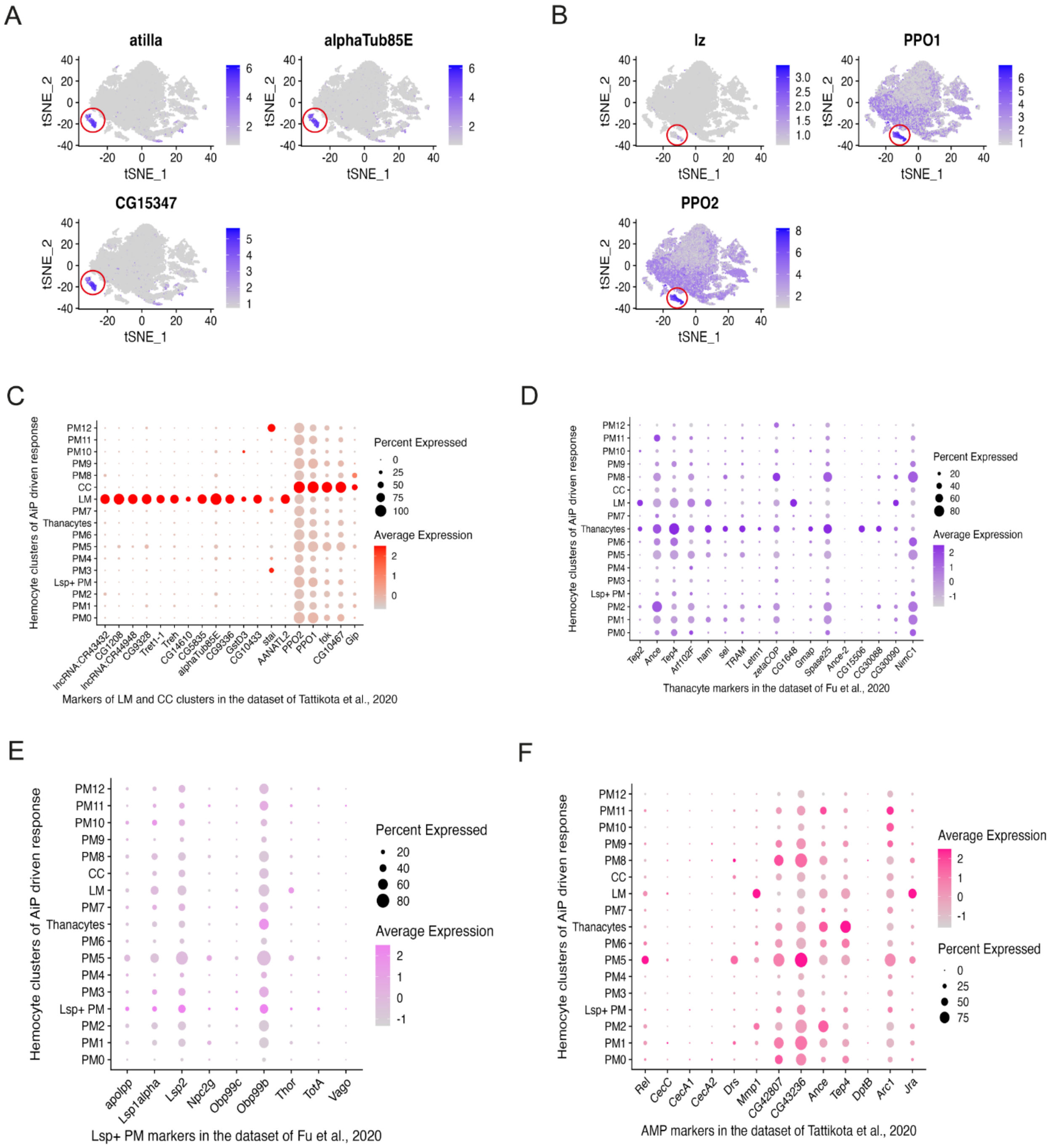

Fig S3: GO Analysis of all new plasmatocyte clusters

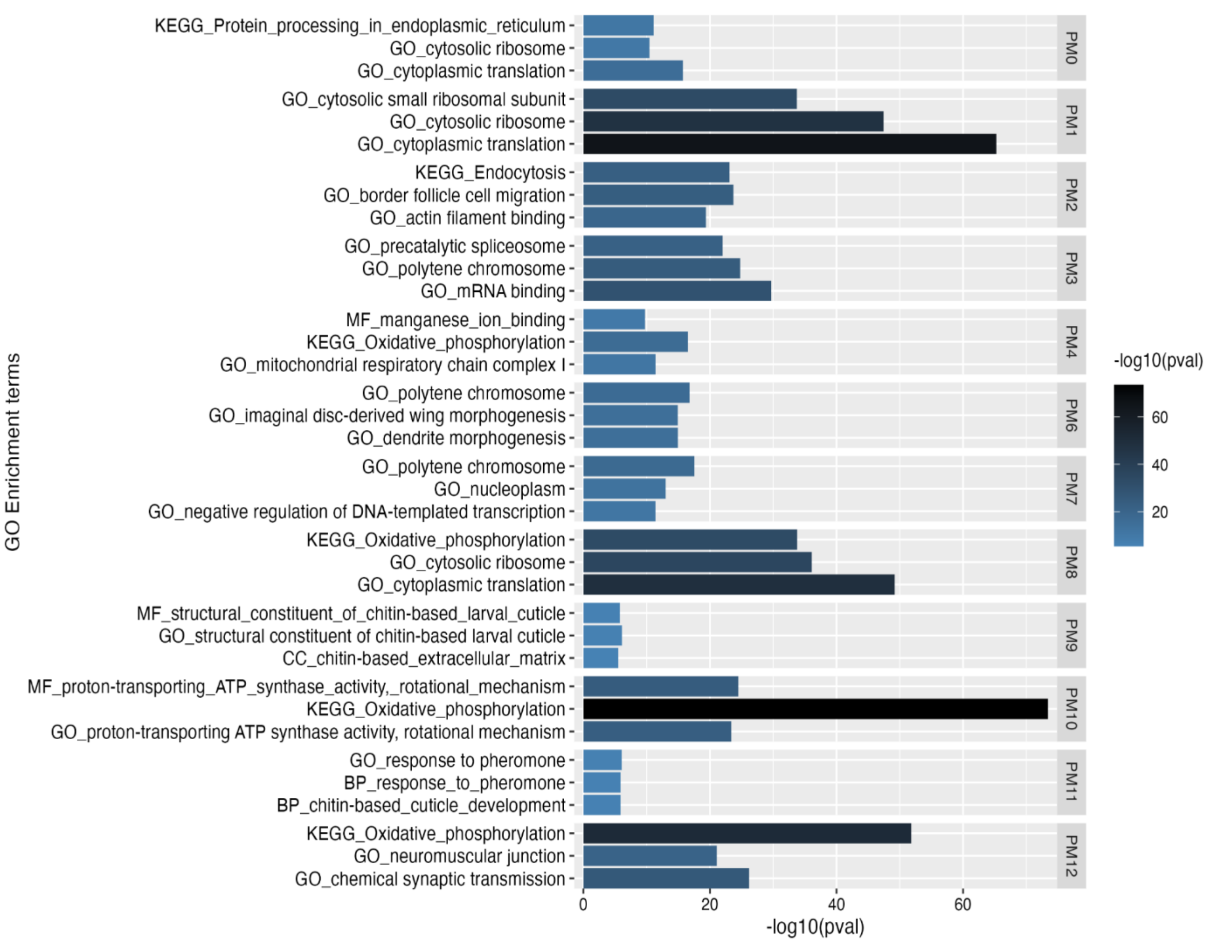
